# Testing the Efforts Model of Simultaneous Interpreting: An ERP Study

**DOI:** 10.1101/212951

**Authors:** Roman Koshkin, Yury Shtyrov, Alex Ossadtchi

## Abstract

We utilized the event-related potential (ERP) technique to study neural activity associated with different levels of working memory (WM) load during simultaneous interpretation (SI) in an ecologically valid setting. The amplitude of N1 and P1 components elicited by task-irrelevant tone probes was significantly modulated as a function of WM load but not the direction of interpretation. Furthermore, the latency of the N1 increased insignificantly with WM load. The P1 latency, however, mostly did not depend on either WM load or direction of interpretation. Larger negativity under lower WM loads suggests deeper processing of the auditory stimuli, providing tentative electrophysiological evidence in support of the Efforts Model of SI. Relationships between the direction of interpretation and median WM load are also discussed.

## I. INTRODUCTION

Unlike in monolingual communication, in simultaneous interpreting (SI) a message in one language is perceived and processed almost concurrently with the production of an equivalent message in another language. To be able to accomplish this feat, besides high proficiency in both the source and target languages, the interpreter must possess a set of specialized skills, including exceptional language switching abilities (Proverbio, Leoni, & Zani, 2004), large working memory (WM) span (Bajo, Padilla, & Padilla, 2000), ability to manipulate WM content and understand incoming discourse while producing a rendering of an earlier portion of the source message in the target language. By its nature, SI is externally paced (Lambert, 1992; Padilla, Bajo, Cañas & Padilla, 1995), indicating the need for cognitive resource management and coping strategies (Chernov, 2004; Kim, 2005; Li, 2010; Liu, Schallert & Carroll, 2004; Mizuno, 2005; Pym, 2008).

In SI an interpreter usually begins interpreting before the speaker has finished a sentence. But the speaker does not wait to move on to the next utterance, regardless of whether the interpreter has completed the translation of the previous chunk (cf. Chernov, 2004; Signorelli & Obler, 2013). Moreover, it may not always be possible or convenient to maintain sequential linearity of the target message relative to the source. For example, interpreters often reverse the order of lists. In some language combinations, e.g. German and English, syntactic constraints force one to wait for the final verb in the German source to construct the target sentence in English (Goldman-Eisler, 1972). Finally, the interpreter may choose to defer translating a word until a good enough equivalent comes to mind, hoping to be able to work it into the target message later. The resulting lag — also referred to as *décalage* or *ear-voice-span (EVS)* in the interpretation studies literature — between the source and the target messages highlights the critical role of WM in the SI pipeline. WM represents a mental space within which to perform the transformations needed for a coherent and accurate target message to emerge.

Under normal circumstances, when the source message is relatively easy to understand and target equivalents are quickly and automatically retrieved from long-term memory (LTM), the interpreter maintains a comfortable décalage, accurately rendering the source message with almost no omissions. But when confronted with a long-winded, dense or obscure passage, the interpreter may be forced out of the comfort zone and temporarily increase the lag to accommodate the need for more time to process it. The lag is similar to debt in that beyond a certain point it becomes difficult to handle. In extreme cases, when the interpreter gets too far behind to speaker, performance quality may be compromised: parts of the source message may get severely distorted or go missing from the translation altogether. This may happen when the interpreter has shifted much of his/her attention away from the current chunk *being* said to finish processing the previous one stored in WM to catch up with the speaker. In sum, large lags are most likely caused by processing difficulties.

On the other hand, when the source message is overall relatively difficult to follow (e.g. when the message is not in the interpreter’s mother tongue), the interpreter may need to allocate extra effort towards understanding. This can be done by shortening the décalage, effectively limiting the amount of information to be processed in working memory. Such a strategy may result in a more literal translation that is likely to be syntactically and grammatically deficient.

In our opinion, the above considerations are best captured by Gile’s Efforts Model (Gile, 1988) which conceptualizes SI in terms of three groups or mental operations, or ‘efforts’: listening, production and memory. Since these efforts are mostly *non-automatic* and *concurrent*, they critically depend on and compete for the limited pool of attentional resources. A major implication of the model is that increased processing demands in one of the efforts can only be met at the expense of another.

To our knowledge, only one study has attempted to test the Effort Model of SI experimentally (Gile, 1999). But as its author himself admitted, “it cannot be said to have led to [its] systematic testing or validation” and also suggested that “precise quantitative measurement” would help to make it more useful. To partially address this concern, in the present paper we used the ERP technique to test one particular prediction of the Efforts Model, namely that increased processing demands on the ‘memory’ effort means less processing capacity available to the ‘listening’ effort. In other words, a higher WM load would create a deficit of attention to the auditory stream. In fact, this hypothesis is quite intuitive but, to our knowledge, has never been tested experimentally in an ecologically valid setting requiring the participants to interpret continuous prose overtly. Electrophysiological evidence supporting it would suggest that interpreters’ brains gate part of the auditory input to be able to properly process the information backlog and reduce the associated processing pressure.

We exploited the previous findings that the P1 and N1 components of the ERP waveform evoked by task-irrelevant probes embedded in a speech stream are larger when the listener is fully focused on the task (cf. Coch, Sanders, & Neville, 2005; Hillyard, Hink, Schwent, & Picton, 1973; Hink & Hillyard, 1976; Teder, Alho, Reinikainen, & Näätänen, 1993; Woldorff & Hillyard, 1991; Woods, Hillyard, & Hansen, 1984). Moreover, a more recent EEG study (Pratt, Willoughby & Swick, 2011) showed that in a multitasking situation — and SI is an extreme case of multitasking (Camayd-Freixas, 2011) — increased WM load decreases attention to the targets. Therefore, the amplitude of these early auditory ERP components can be used as a suitable and temporally precise index of interpreters’ attention to the spoken *source* message.

Our assumption that WM overload reduces attention to the auditory stream, which in turn modulates the ERP waveform, aligns well with the evidence that both WM and attention may utilize a common pool of neural resources (Sabri et al., 2014). As demonstrated by fMRI studies, attention and WM are subserved by largely the same brain areas (Awh, Vogel & Oh, 2006; Awh & Jonides, 2001; Corbetta & Shulman, 2002; Cowan, 2000; Mayer et al., 2007; Ungerleider, 2000; Zanto, Rubens, Thangavel & Gazzaley, 2011).

We also examined whether the direction of interpretation has any effect on the early ERP components and sought to identify its potential interaction with WM load. We were motivated by an informal survey conducted prior to this study. In it, out of 32 professional simultaneous interpreters, 29 reported that, all else being equal, interpreting from L2 into L1 was much more difficult than in the opposite direction. This is at odds with the findings of a PET study (Rinne et al., 2000) on Finnish L1 interpreters that suggested L1-L2 interpretation was more difficult. The interpreters we surveyed also said the most difficult part for them was to *understand* the source message in L2—and understanding is part of the *listening* effort according to Gile (1988, 1995, 1999). Based on these observations and the prediction of the Efforts Model we predicted that this subjective difficulty would result in a significant difference in median WM loads. Finally, we looked into a possible connection between the direction of interpretation and attention as indexed by the early ERPs.

Achieving the goals of the study required defining a method of estimating WM load. The most obvious and straightforward approach to do it would be to simply assume that WM load is proportional to the number of words that an interpreter lags behind the source message at any given time. In the time intervening between the moment of hearing a word and finishing its translation overtly, the interpreter has to store it in WM at least until it is further processed as appropriate to the situational context. If we assume that the cognitive effort it takes to process the most frequent function words—especially articles and prepositions—is negligible, we can exclude them from WM load estimation. However, this approach would be an oversimplification. First, it is predicated on the assumption that each word strains WM capacity equally. More frequent words must have stronger mental representations and a larger network of associations than relatively rare ones. In what is known as the *word frequency effect* (Gardner, Rothkopf, Lapan & Lafferty, 1987; Rubenstein, Lewis, & Rubenstein 1971; Scarborough, Cortese & Scarborough, 1977) more frequent words are processed faster. Therefore, they must be translated more easily and cleared from working memory relatively quickly to make room for processing the next chunk of the source message. Conversely, words that are less familiar take more time and effort to process, which delays the retrieval of equivalents from LTM and increases WM load. Second, such an approach does not take into account that words in the source message are never chunked together such that they are stored in WM as a single mental object. Third, it assumes that WM load is reduced at the moment when a word has been translated overtly. This is not always the case because interpreters continue to post-process their own translation, in a phenomenon known as self-monitoring (Gerver, 1974, 1975; Gile, 1995, 2008; Setton, 1999). This means that after a word (or, more generally, a chunk) is translated overtly, it is not cleared from WM immediately, but at a later time, when the interpreter is satisfied with his or her translation. An opposite situation may happen—and this is the fourth reason—when the interpreter can use informational redundancy of the source message to predict an idea *before* it is fully uttered by the speaker (Chernov, 2004). For example, if the source message has multiple references to the United Nations Organization, it is easy to finish the translation before the offset of the phrase *the United Nations Organization*, in other words get ahead of the speaker. The above considerations mean that WM load estimates will be inherently noisy. To mitigate this error of measurement, we sought to maximize the sample and explored several alternative methods of WM load estimation, which we describe below.

## II. METHOD

### Participants

Nine males (aged 25-47, *M* = 36.9, *SD* = 6.757) participated in the study. All were qualified interpreters holding a university degree in translation and interpreting, L1 Russian speakers, with an average of 10.65 (*SD* = 6.675) years of professional SI experience. None of them reported substance abuse or a neurological disorder. All signed an informed consent form.

### Procedure and materials

The participants were asked to interpret 8 speeches (4 Russian and 4 English) from the 6849^th^ United Nations Security Council Meeting. The discussions at the meeting focused on the rule of law, a topic unlikely to elicit an emotional response. Our expectation was that the UN terminology would be familiar enough to the participants so that they could deliver quality translation with little or no preparation. All the speeches had been originally presented in the speaker’s mother tongue (2 in French, 6 in Spanish) with simultaneous interpretation into all the six official UN languages, including Russian and English, in line with the standard UN practice. We deliberately chose speeches originally delivered in a language other than Russian or English. This was necessary to avoid a potential bias due to some subtle properties (e.g. idiosyncratic syntax) that may have been present in the translated, but not in the original texts, and vice versa. To control for a potentially confounding effect of varying delivery speed, we had the written Russian and English translations read at a slow constant rate by a bilingual speaker (female) highly proficient in both Russian and English. After the recording we used Audacity (http://audacityteam.org), open audio editing software, to adjust the recording to ensure a constant delivery rate of 105 words per minute (wpm). This was an appropriate measure since if these texts had been read out loud within the timeframe of the original speeches, the text-to-text variance of the reader’s rate of speech would have had been quite high (*M* = 121.75 wpm, *SD* = 30.64 wpm for Russian, *M* = 125 wpm, *SD* = 42.1 wpm for English). Thus we eliminated the possible effects of individual speaker’s voice features such as rate, pitch, timbre, loudness, prosody and accent. The total playback time was 53 minutes (excluding periods of rest between the speeches).

The audio streams and the task-irrelevant probe stimuli were played by a custom script running under PsychoPy (Peirce, 2008). The probes were 440-Hz 52-ms pure sine tones (including a rise and fall period of 4 ms) delivered with a jittered inter-stimulus interval (ISI) of 450–750 ms (M = 600 ms). These parameters were selected empirically to maximize the number of probes per second of experimental time while minimizing the effect of diminished ERP amplitude with shorter ISIs (Teder et al., 1993).

To control for order effects, the speeches were delivered to the participants in a pseudo-random fashion according to a Latin square such that for every participant the texts’ order was different.

The texts were played through earbud headphones (Sennheiser MX 170, 32Ω, Germany). Before the experiment, the participants did two 30-second practice runs, in which they were asked to adjust the volume to a comfortable level and interpret an excerpt from a speech delivered at the same UNSC session, but not included in the experimental material.

Participants were seated comfortably in a reclining chair in an electrically and acoustically shielded room. To reduce oculo-motor and muscle artifacts, they were instructed to sit still, relax, minimize eye movements and articulate their translation as quietly as possible. The translation was recorded using a Boya BY-M1 capacitor microphone for subsequent offline transcription and timecoding.

### Working memory load estimation

The number of content words in the source-target provides a measure of instantaneous WM load. However, a potentially more precise estimate could be achieved by scaling every word in the source texts by its min-max normalized log-transformed frequency, 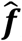, in a representative language corpus. Min-max normalization maps a word’s log-transformed absolute frequency in the corpus, ***f***, to a value in the range between 0 and 1, while log-transformation helps ensure a better fit for the linear model describing the relationship between a word’s rank and frequency in the corpus. For example, the WM load associated with the word ‘peace’, 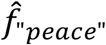, after controlling for its frequency in the corpus is given by:

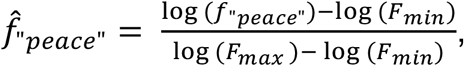

where *F_max_* and *F_min_* are the absolute frequencies, respectively, of the most and the least frequent word in a given corpus, and *f_“peace”_* is the absolute frequency of the word ‘peace’ in the corpus. Estimating WM load this way seems appropriate since word frequency is a major factor determining word recognition (Burgess & Livesay, 1998). More frequent words elicit quicker responses than less frequent ones (Dobbs, Friedman, & Lloyd, 1985; Gernsbacher, 1984). In fact, Whaley (1978) argued that word frequency was “by far the most powerful predictor” in a lexical decision task. For Russian source texts we chose the ruTenTen (14.5 billion words) and the British National Corpus (112.2 million words) accessed through Sketchengine (https://www.sketchengine.co.uk/). Before looking up a word’s frequency in the corpus, we used the NLTK package (Bird, Klein, & Loper, 2009) to lemmatize each word.

The first method described above is based on the number of content words. However, it can be extended by scaling each word according to its syllabic length, a potentially important factor influencing the speed of processing. This extension makes sense since the phonological loop — an essential component of auditory WM (Baddeley, 1992; Baddeley & Hitch, 1974) — has a capacity of about 2 s unless the information stored in it is constantly rehearsed and/or processed.

In summary, we assessed the performance of three different methods to estimate WM load, specifically in the number of (a) content words^1^ (CW); (b) all the words (both function and content) weighted by their frequency in the language (CL), (c) content words weighted by their respective syllabic length (SYL). Changes in WM load were assumed to occur at the time of word offset. Since the probes were delivered at random intervals not corresponding to word offsets, we used linear interpolation to estimate WM load at probe onsets (see Figure 1).

**Figure 1.**
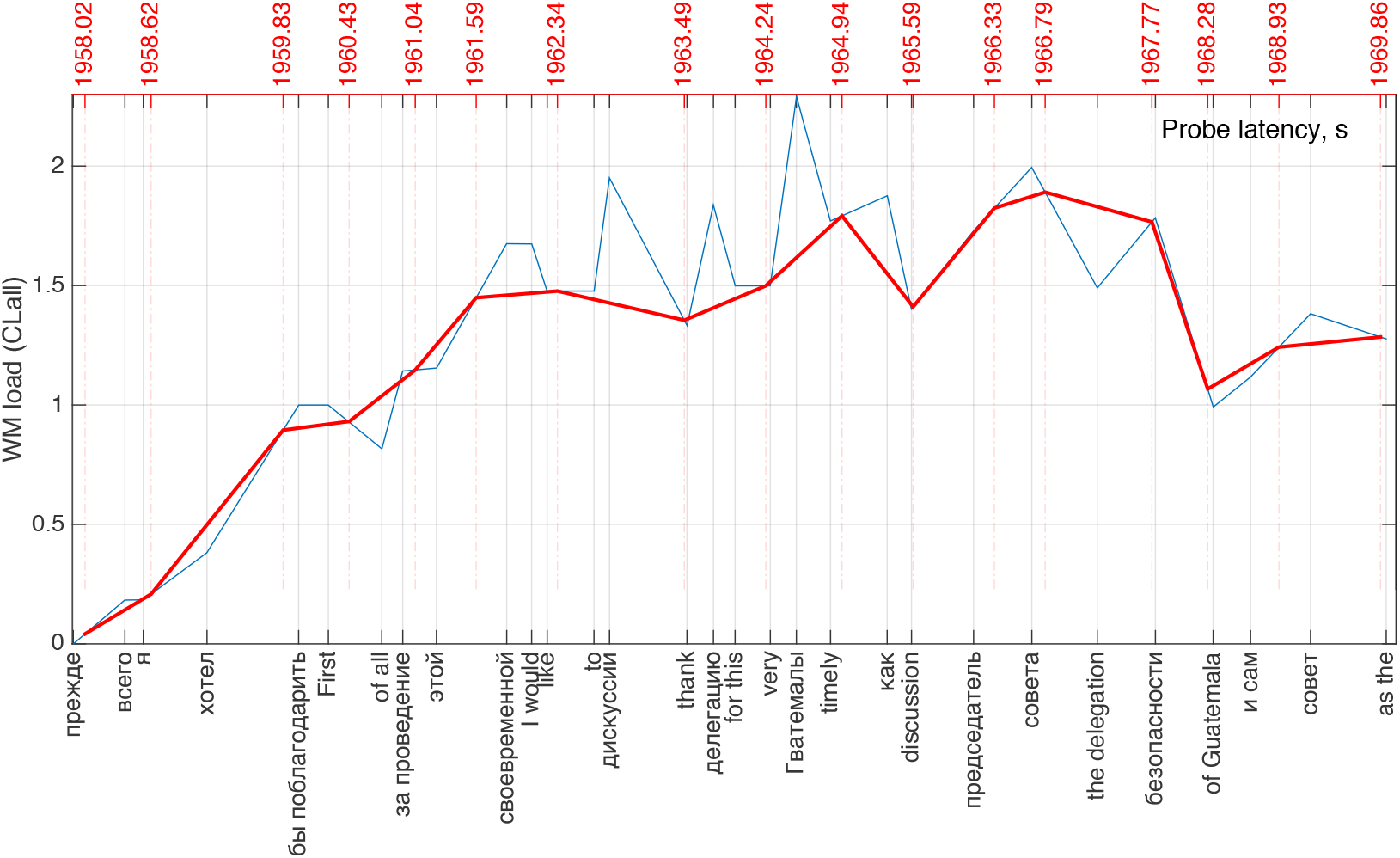
A slice of a time-coded transcript showing WM load dynamics over time. The blue line represents WM load at word offsets. The red line captures linearly interpolated values of WM load at probe onsets. Red numbers along the top axis show probe onset times.

### Transcript time coding and WM load estimation

After all the participants’ EEG data were recorded, we made transcripts of both the original and translation. These were manually time-stamped and reformatted to allow us to calculate the different WM load estimates as described above. This work was partially automated using several custom functions in VBA running under Microsoft Excel. Figure 2 illustrates the resulting data.

**Figure 2.**
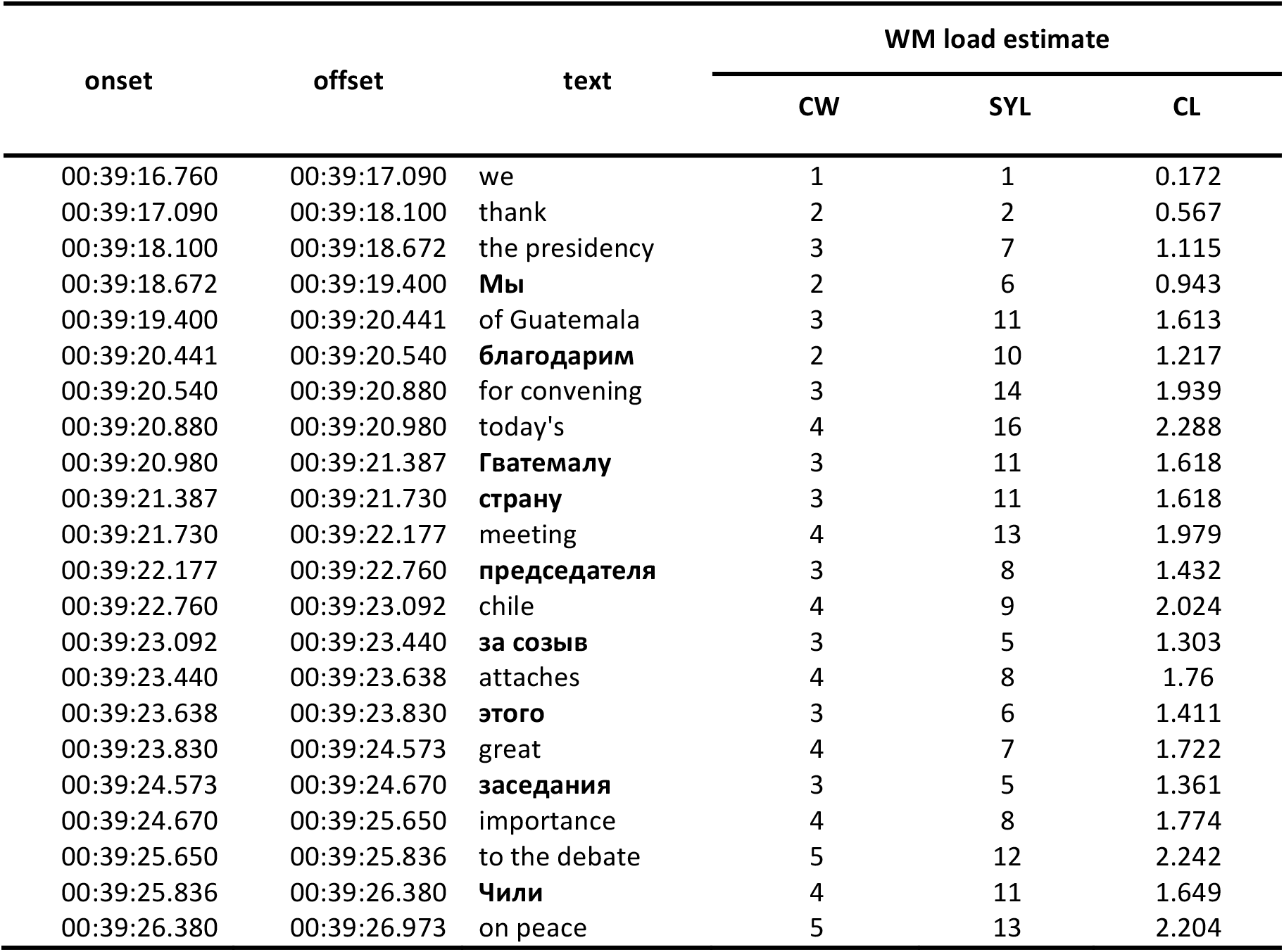
Illustration of a time-coded transcript with WM load estimates.

### EEG data acquisition and pre-processing

The electroencephalogram (EEG) was continuously recorded using the ActiCHamp recording system and BrainVision PyCorder software (Brain Products GmbH) at the sampling rate of 2000 Hz from 32 scalp sites with active electrodes seated in an elastic cap.

EEG preprocessing and analysis were carried out using the EEGlab toolbox (Delorme & Makeig, 2004) for MATLAB (The Mathworks, Natick). Raw datasets were downsampled offline to 250 Hz, converted to a linked-mastoids reference (TP9 and TP10) and bandpass-filtered from 0.1 to 30 Hz (6 db/oct) using a zero-phase FIR filter. At the next step we ran independent component analysis (ICA) using the *binica* algorithm in EEGlab to remove the two most dominant independent components corresponding to oculomotor artifacts. Then, to address the issue of muscle noise produced by constant articulation in the experiment, we cleaned continuous data with the artifact subspace reconstruction (ASR) algorithm (Mullen et al., 2015) with the burst parameter set to 4 standard deviations^2^.

To ensure the possibility of averaging the data epochs within specific WM load (*low, medium* or *high*), we changed event codes in the EEG dataset to reflect WM load at the probe onset latency estimated with the five different methods described above.

The continuous EEG was then chunked into epochs of 500 ms post-stimulus onset including a 100 ms pre-stimulus baseline, yielding about 5840 epochs per subject. The epochs were screened for any residual artifacts that may have survived the ASR- and ICA-based cleaning stage. Then we averaged the epochs within direction of interpretation and WM load estimated at time zero, i.e. the moment of probe onset, using each of the five methods.

### Data Analysis

After cleaning the EEG datasets, we performed several statistical tests of our hypotheses. First, we wanted to check for possible main effects (and interaction) of WM load and interpretation direction on the average ERP amplitude in the N1 and P1 range. The P1 and N1 component amplitudes and latencies were computed for 40-ms windows around the observed P1 and N1 grand average peaks (40-80 and 120-160 ms, respectively, post stimulus onset). We used a 4-way repeated measures analysis of variance (ANOVA) with Direction (L1->L2, L2->L1), WM Load (Low, Medium, High), Anteriority (front, center, back) and Laterality (left, middle, right) as factors. The Kolmogorov-Smirnov test of the mean P1 (D = 0.047213, *p* < 2.2e-16 P1) and N1 (D = 0.041403, *p* < 2.2e-16 N1) indicated a violation of normality. To address this, we used the Johnson transformation^3^ (after the transformation *D* = 0.0037101, *p* = 0.121 for P1; *D* = 0.0032198, *p* = 0.2799 for N1). Unless stated otherwise, all the statistical tests were done in R (R Development Core Team 2016) using the *ez* library.

## III. RESULTS

### WM load estimates from behavioral data

Figure 3 shows the distributions of WM loads by participant and direction of interpretation. We first consider the distributions of WM load estimated by the most straightforward methods CW and SYL described above. All the participants’ median WM loads estimated with the CW and SYL methods were smaller in the L2>L1 than in the opposite direction. A Wilcoxon^4^ signed rank test showed this difference to be significant: *V* = 45, *p* = 0.003906. However, the difference between WM loads calculated using the other method (CL) did not reach statistical significance:, *V* = 6, *p* = 0.05469.

**Figure 3.**
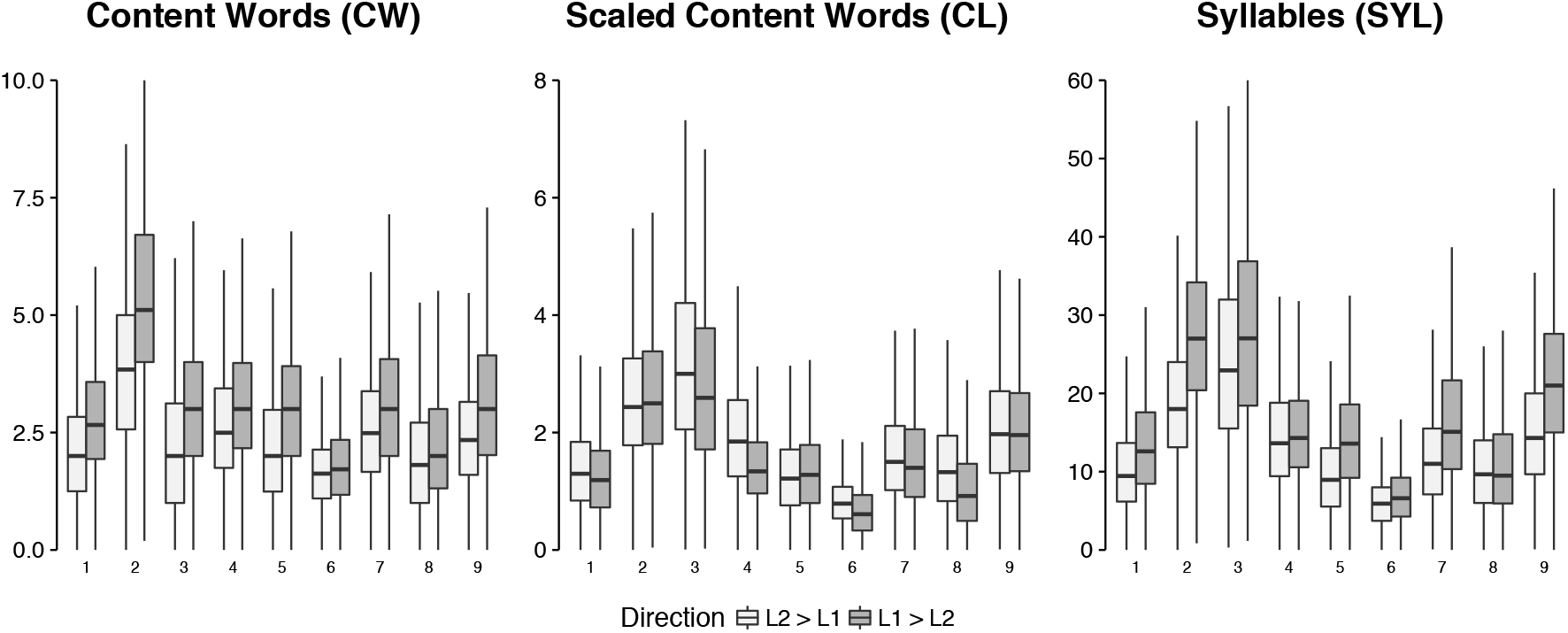
Distributions of WM load estimates by subject and direction of translation calculated using the three different methods.

### EEG Results

To obtain an equal number of observations within each cell of our fully crossed experimental design, the boundaries between the high, medium and low WM load conditions should have been set at the 33^rd^ and 66^th^ quantiles of each subject’s WM load distributions (both for English-Russian and Russian-English). However, as can be inferred from Figure 3, there is a certain ‘comfort zone’ that interpreters keep to (or are forced to keep by the very conditions of the interpretation task) most of the time. Therefore, splitting the epochs into three equal WM load groups is not likely to capture a potential effect on the neural activity due to the associated WM load. With that in mind, we labeled an epoch as *low* if the WM load at the probe onset was below the 10th quantile, *medium* if the WM load fell between the 10^th^ and 90^th^ quantile, and *high* if it exceeded the 90^th^ quantile in the corresponding WM load distribution. Although such a choice may seem arbitrary, it was a tradeoff between maximizing the effect size and sacrificing statistical power due to inflated error in the *low* and *high* WM condition. Mindful of large between-subject and between-language variance in the WM load distributions, we calculated the boundary quantiles for each participant and direction of translation individually obtaining 18 pairs (9 participants x 2 source language) of subject- and direction-specific condition boundaries.

Figure 4 shows grand average ERPs at the vertex electrode (Cz) elicited by the probes as a function of WM load.

**Figure 4.**
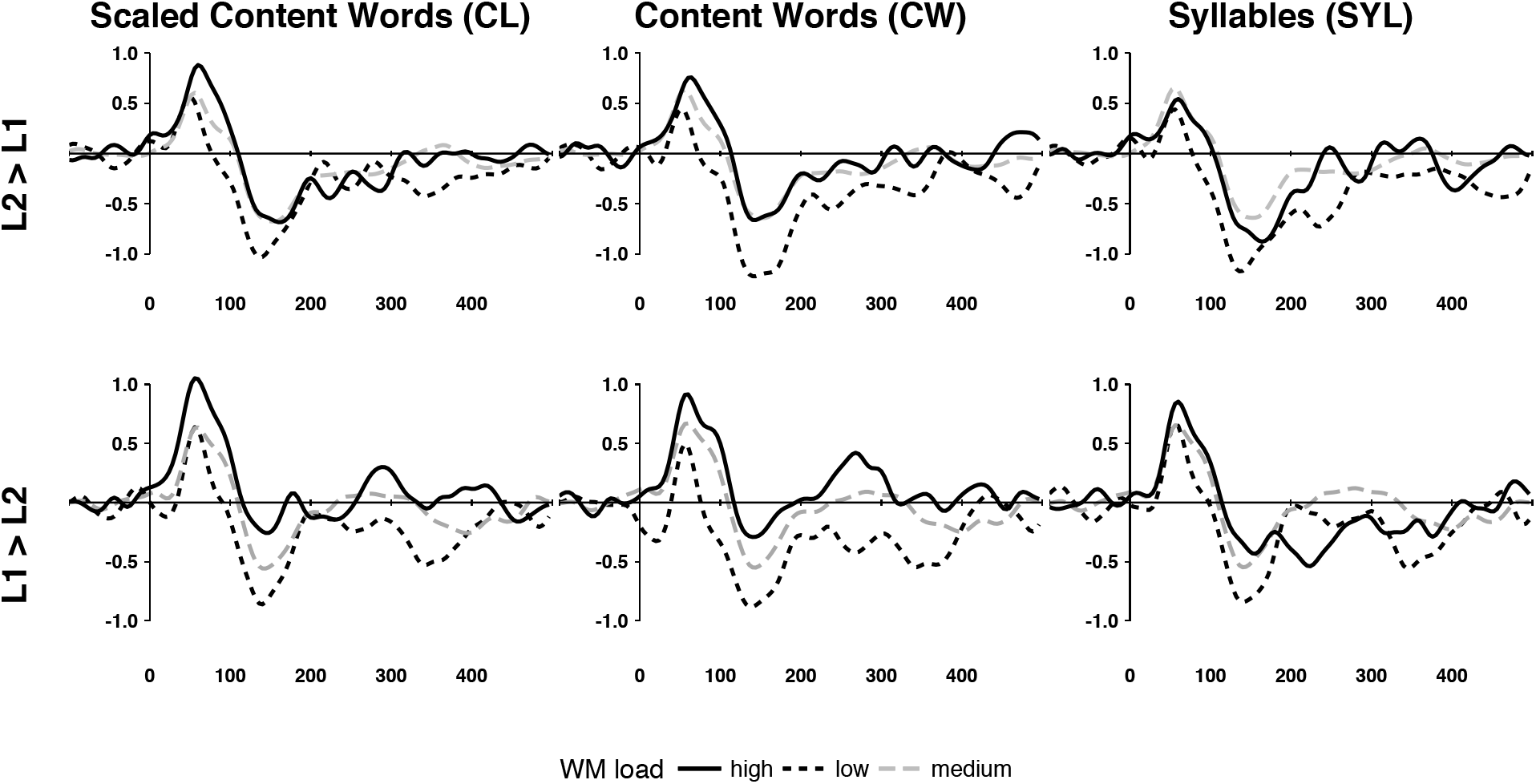
Grand average ERPs for L1 > L2 and L2 > L1 direction at Cz. The ERPs are plotted for the three different methods of estimating WM load. Y-axes display microvolts and X-axes milliseconds.

#### P1 amplitude analyses

Regardless of the WM load estimation method used, we found neither a main effect of interpretation direction (CL [*F*(1, 8) = 0.17, *p* = .690, 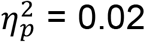]; CW [*F*(1, 8) = 0.39, *p* = .551, 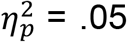]; SYL [*F*(1, 8) = 0.09, *p* = .915, 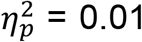]), nor WM Load × Direction interaction (CL [*F*(1.08, 8.66) = 0.12, *p* = .753, 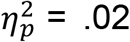]; CW [*F*(2, 16) = 0.55, *p* = .585, 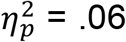]; or SYL [*F*(1.11, 8.91) = 0.49, *p* = .520, 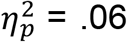]). The main effect of WM load reached significance for the two methods based on the number of content words in the source-target lag (CL [*F*(1.24, 9.92) = 5.43, *p* = .037, 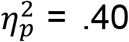]; CW [*F*(2, 16) = 3.89, *p* = .042, 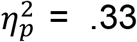], but not the one based on syllables (SYL [*F*(2, 16) = 0.09, *p* = .915, 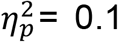]). Although the effect of WM load was stronger over the frontal sites, the Anteriority × Load interaction reached significance only for the CL method [*F*(2.25, 18.0) = 7.83, *p* = .003, 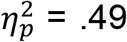], but not for the other methods (CW [*F*(1.29, 10.33) = 1.83, *p* = .208, 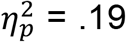; SYL [*F*(1.33, 10.67) = 0.08, *p* = .846, 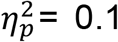]).

#### N1 amplitude analyses

Similarly to the P1 range, we found neither main effect of interpretation direction (CL [*F*(1, 8) = 0.01, *p* = .914, 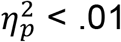]; CW [*F*(1, 8) = 0.35, *p* = .570, 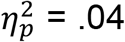]; or SYL [*F*(1, 8) = 1.21, *p* = .302, 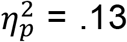]), nor WM Load × Direction interaction (CL [*F*(2, 16) = 0.05, *p* = .952, 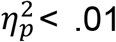]; CW [*F*(2,16) = 0.08, *p* = .925, 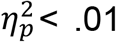]; or SYL [*F*(1.14, 9.16) = 0.20, *p* = .697, 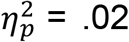]). However, the main effect of WM load was consistently significant for all the WM load estimation methods (CL [*F*(2, 16) = 8.06, *p* = .004, 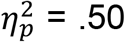]; CW [*F*(2, 16) = 9.87, *p* = .002, 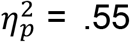]; or SYL [*F*(2, 16) = 7.88, *p* = .004, 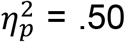]). As in the P1 range, the effect of WM load was more pronounced over the frontal sites. However, the Anteriority × Load interaction reached significance only for the CW method [*F*(2.00, 16.1) = 3.87, *p* = .043, 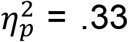], but not for the other methods (CL [*F*(1.72, 13.74) = 3.02, *p* = .087, 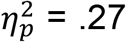; SYL [*F*(1.69, 13.56) = 2.20, *p* = .153, 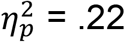]).

Figures 4 and 5 show grand average ERP amplitudes at the Cz channel as a function of WM load.

**Figure 5.**
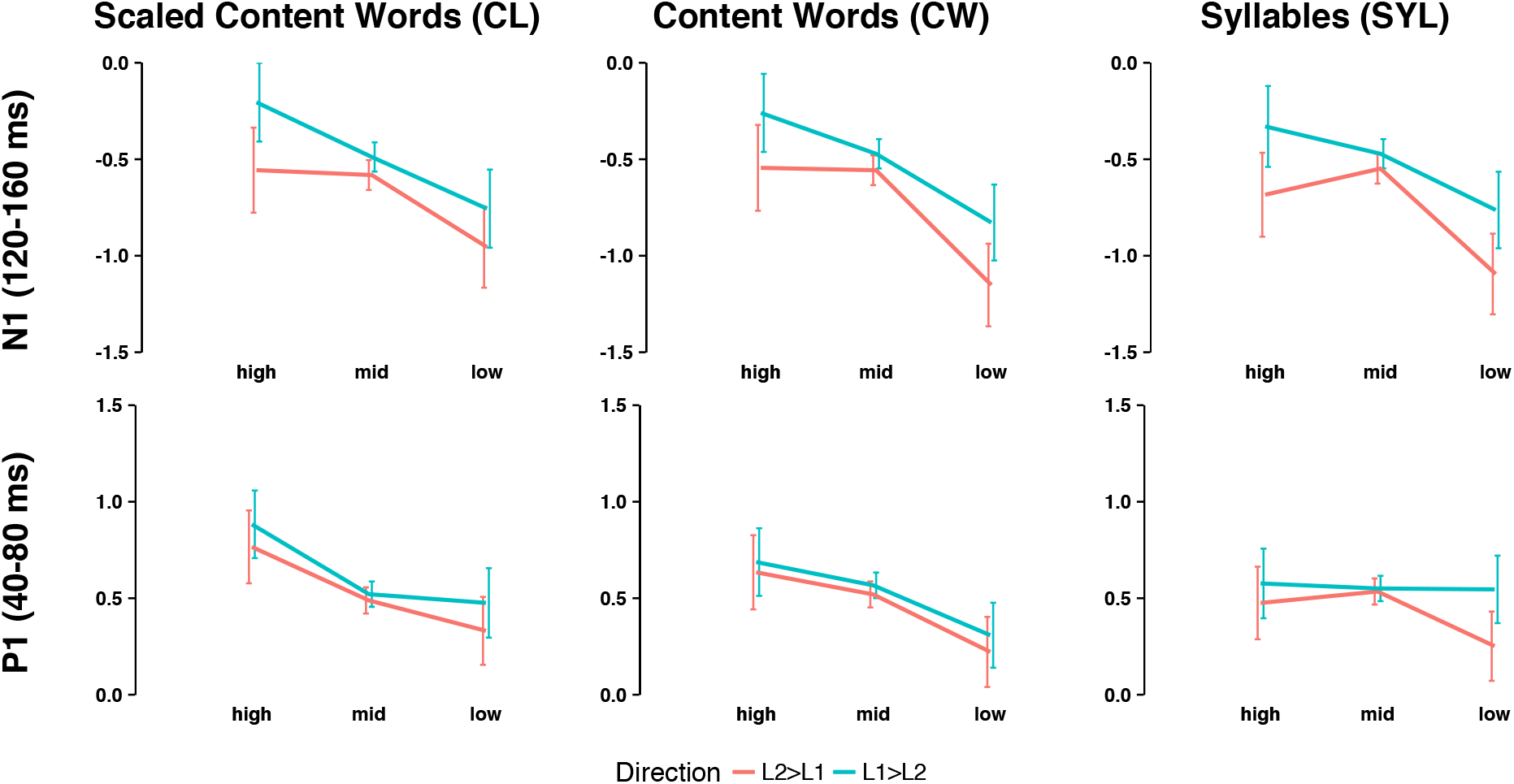
Grand average P1 and N1 amplitudes at Cz as a function of WM load. Vertical bars represent standard errors of the mean.

Figure 6 shows the scalp activations computed on grand average ERP data. It is clearly seen that the P1 peak (~60 ms post stimulus onset) was topographically centered around frontal midline electrodes (Fz and Cz), while the N1 peak is slightly offset to the right.

**Figure 6.**
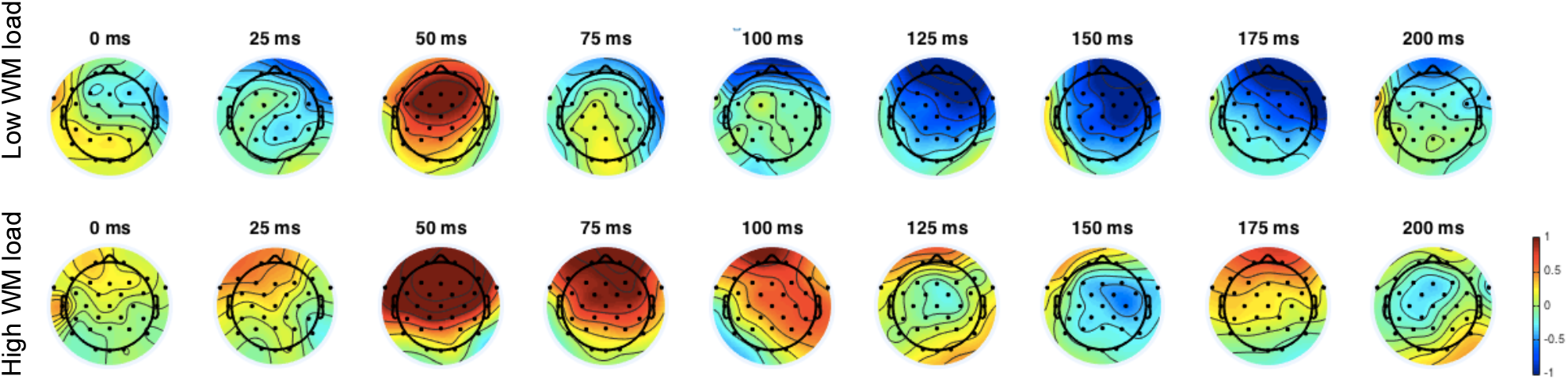
Grand average ERPs scalp maps.

#### Peak latency analyses

Peak latencies for visually identifiable ERP components at the Cz channel, specifically P1 and N1, are reported in Table 1. The p-values for the difference between the peak latencies were calculated using a permutation test with 5000 resamples.

**Table 1.** Peak latencies in milliseconds post stimulus onset in the P1 and N1 range for the low and high WM load conditions at the Cz electrode. The estimated permutation p-values are reported for the latency differences between the high and low WM load.

| Russian - English |  |  |  |  |  |  |
| --- | --- | --- | --- | --- | --- | --- |
| WM load estimation method | P1 (40-80 ms) |  |  | N1 (120-160 ms) |  |  |
|  | Low WM load | High WM load | p | Low WM load | High WM load | p |
| CL | 52 | 56 | 0.0014* | 140 | 168 | 0.2157 |
| CW | 56 | 60 | 0.0257* | 140 | 148 | 0.8600 |
| SYL | 56 | 56 | 0.4886 | 140 | 168 | 0.2814 |

| English – Russian |  |  |  |  |  |  |
| --- | --- | --- | --- | --- | --- | --- |
| WM load estimation method | P1 (40-80 ms) |  |  | N1 (120-160 ms) |  |  |
|  | low WM load | High WM load | p | Low WM load | High WM load | p |
| CL | 56 | 56 | 0.2142 | 136 | 148 | 0.2342 |
| CW | 56 | 56 | 0.1771 | 136 | 148 | 0.4185 |
| SYL | 56 | 56 | 0.1042 | 136 | 152 | 0.3400 |

The data in Table 1 suggest that peak N1 latencies in the high WM load condition tend to be greater (and are never smaller) than in the low WM load condition. However, this trend is very weak, especially for the P1 peak latency, and does not consistently reach significance for all the WM load estimation methods.

## IV. DISCUSSION

The present study aimed to find electrophysiological evidence in support of the Efforts Model (Gile, 1988, 1995, 1999) which predicts, in particular, that WM overload during simultaneous language interpretation should decrease the amount of processing capacity available to the ‘listening effort’ and therefore affect the processing of auditory stimuli. Although WM load can be conveniently estimated if we assume it to be proportional to the size of lag between the source and the target (in number of content words), getting a precise behavioral measure of attention during SI would have been problematic. However, as was previously demonstrated in non-interpreters, the amplitude of early ERP components time-locked to task-irrelevant probe stimuli could be used as an electrophysiological index of attention. Given that and based on the Efforts Model, we expected larger negativity in the N1 range at small WM loads.

Consistent with our hypothesis, we found that the amplitude of the N1 component elicited by the task-irrelevant pure tone probes was more negative at lower WM loads. Conversely, the N1 amplitude was less negative at higher WM loads suggesting that the brain temporarily attenuates or suspends the processing of auditory stimuli to more efficiently process and manipulate WM backlog, reduce the lag and cognitive load. The question whether this is controlled consciously (i.e. is a choice based on strategic judgement), or automatically (i.e. is a skill acquired though training and used unconsciously) is treated elsewhere (cf. Fukuii and Asano 1961, Oléron and Nanpon 1965, Kade and Cartellieri 1971, Daró and Fabbro 1994, Osaka 1994, Padilla 1995, Gran and Bellini 1996, Chincotta and Underwood 1998). A correlational study on a larger sample of interpreters with different levels of experience may in the future help answer the question with more certainty.

A visual inspection of the ERP waveforms revealed that the effect of WM load in the P1-N1 range may not have been due only to the enhanced negativity, but also to decreased latency of the P1 and N1 peaks for low WM load. Fehér, Folyi & Horváth (2011) reported a shorter N1 latency for attended stimuli, which reinforces the case for attention acting as a latent variable mediating the effect of WM load on the N1. Although our results did suggest greater N1 latencies at larger WM load, they were not statistically significant.

As part of our second research question, we wished to establish if the subjectively greater difficulty of SI from L2 into L1 reported by our informal respondents, was due to direction-specific WM load management differences and, if any, to find their electrophysiological correlates. The behavioral data showed a pattern of smaller median WM loads in the L2-L1 direction, which seems to be consistent with the predictions of the Efforts Model. However, further interpretation of this pattern is difficult due to no significant main effect of direction on the ERPs (P1/N1) or interaction between direction and WM load.

The fact that the two interpretation directions had significantly different median WM loads may be due to several reasons. First, because L2 words on average are less ‘familiar’ than L1 words, they have weaker mental representations than the corresponding L1 words. As a result, L2 words take more time to react to in lexical decision tasks (Gernsbacher, 1984). For our participants (Russian L1, English L2 speakers) this indicates that the perception and analysis of L2 speech is more difficult than L1. Moreover, this difficulty is not offset by the relative ease of producing a meaningful target message in the mother tongue, which is evidenced by shorter lexical retrieval times for L1 words (cf. Schoonen et al., 2003). Secondly, WM in L2 is known to be slower and less efficient than in L1 (Ardila, 2003). Perhaps, when working from L2 into L1, interpreters conserve processing capacity by reducing WM load, while in the opposite direction they can afford to keep in WM more L1 than L2 content words. Thirdly, interpreting from Russian into English—again, all else being equal—appears less of a challenge because Russian words are on average syllabically longer than in English (Gurin, 2009). Thus, a source message in Russian is informationally more sparse and easier to process than a comparable English message delivered at the same rate measured in syllables per unit time. Conversely, in Russian-English interpretation the target English discourse is syllabically more compact and therefore should cause less interference due to acoustic overlap with the source message. In other words, if we assume word-for-word rendering, the interpreter has less time to ‘unpack’ the English message: English words are shorter than Russian requiring more time to complete the articulation of the corresponding Russian translation. Not only does this create greater and potentially disrupting acoustic interference with the process of listening to the English source, it is possible that attention is engaged longer in the self-monitoring stage of the interpretation cycle. Further research is needed to address this question which can use different language combinations to vary the above parameters systematically. However, at least one study (Pellegrino, Coupé & Marsico, 2011) based on seven languages—unfortunately, not including Russian—found that in spoken communication the average amount of information transferred per unit time is subject to substantial, however statistically insignificant, cross-language variation. It is possible that the obvious difference in WM loads as a function of source language is due to different information densities of the languages used here.

Although it is impossible to generalize our results to situations with any other source-target language pair, our data suggest that the early sensory processing does not depend on whether the source is in the mother tongue (L1) or a foreign language (L2). We can argue, at least, that the main source of subjective difficulty in the English-Russian interpreting does not lie in the perceptual stage of processing.

It would be interesting to know if our pattern of results will replicate in language pairs other than those employed here. Languages may have vastly different syntaxes and grammars, which means that interpreters have to change their strategies depending on the direction of interpretation. For example, when working from German, interpreters are forced to increase their décalage (Goldman-Eisler, 1972) waiting for the predicate which often appears at the end of the clause, which means larger WM loads overall.

On a practical note, our results justify a recommendation to keep WM load within reasonable limits during SI. The question about how this can be achieved is very relevant. Several studies (Bartłomiejczyk, 2006; Chernov, 1994; Li, 2010) have shown that simultaneous interpreters use a range of strategies to manage their processing load (e.g. omitting redundancies in the source speech). None of them, however, have attempted to identify the neural states that determine—or at least bias—the choice of a particular strategy.

Further neuroimaging studies of SI are needed to validate and/or refine theoretical models of SI and will help to guide interpreters, SI instructors and the interpretation industry towards better interpretation practices, more efficient curricula and higher standards of service.

## VII. AUTHOR CONTRIBUTIONS

RK, AO and YS designed the research. RK conducted the research, performed analyses and drafted the manuscript. YS and AO contributed to the analysis and interpretation of the results and the manuscript preparation. All authors agreed on the final version of the manuscript.

## VIII. ACKNOWLEDGMENTS

The study was funded by the Russian Academic Excellence Project ‘5–100’.

## Footnotes

1 In this study we used the following definition of a function word: articles, prepositions, auxiliary

2 According the authors of the ASR algorithm implementation, 4 sigma ensures mild cleaning.

3 Implemented in the *jtrans* library for R.

4 Since all the WM load distributions were clearly right-skewed, a non-parametric test was in order.

